# Active presynaptic ribosomes in mammalian brain nerve terminals, and increased transmitter release after protein synthesis inhibition

**DOI:** 10.1101/295543

**Authors:** Matthew S. Scarnati, Rahul Kataria, Mohana Biswas, Kenneth G. Paradiso

## Abstract

Presynaptic neuronal activity requires the localization of thousands of proteins that are typically synthesized in the soma and transported to nerve terminals. Local translation for some dendritic proteins occurs, but local translation in mammalian presynaptic nerve terminals is difficult to demonstrate. Here, we present evidence for local presynaptic protein synthesis in the mammalian brain at a glutamatergic nerve terminal. We show an essential ribosomal component, 5.8s rRNA, in terminals. We also show active translation in nerve terminals, in situ, in brain slices demonstrating ongoing presynaptic protein synthesis. After inhibiting translation for ~1 hour, the presynaptic terminal exhibits increased spontaneous release, and increased evoked release with an increase in vesicle recycling during stimulation trains. Postsynaptic response, shape and amplitude were not affected. We conclude that ongoing protein synthesis limits vesicle release at the nerve terminal which reduces the need for presynaptic vesicle replenishment, thus conserving energy required for maintaining synaptic transmission.

## Introduction

Synaptic transmission requires the synthesis, localization, interaction and ongoing replenishment of thousands of pre- and postsynaptic proteins (Witzmann, et al., 2005; Gonzalez-Lozano, et al., 2016; Loh, et al., 2016). The location and stoichiometry of each protein is highly regulated to maintain the necessary levels of precision and fidelity of signaling across the synapse. The highly structured and polarized morphology of neurons, with axons and dendrites that can project long distances, creates a unique challenge to maintain sufficient levels of numerous necessary proteins at distant locations (Alvarez, et al., 2000; Maday, et al., 2014; Tasdemir-Yilmaz and Segal, 2016). In addition, these remote regions need to rapidly modify the magnitude and duration of their responses, which can require changes in pre- and postsynaptic protein expression levels. Synaptic proteins are typically thought to be synthesized in the soma and transported to synapses, but several groups have demonstrated that some postsynaptic proteins can be synthesized locally in dendrites (Pfeiffer and Huber, 2006; Jung, et al., 2014; Rangaraju, et al., 2017). Over the past decade, RNA based mechanisms have been discovered that respond to extrinsic signals that affect postsynaptic local translation in dendrites to modify activity at specific regions (Liu-Yesucevitz, et al., 2011; Yoon, et al., 2016). This is possible due to the targeting of coding and non-coding RNA (Vo, et al., 2010) with RNA binding proteins, and the presence of ribosomes that are located in, or moved to specific neuronal regions or compartments (Ostroff, et al., 2002). This allows the neuron to have the necessary components in place to translate specific dendritic proteins on-site, in response to specific signals. The role of local translation in resting and sustained levels of synaptic transmission is a major issue of interest.

Local protein synthesis is thought to provide a faster and more efficient mechanism for neurons to maintain or alter activity levels and respond to rapidly changing inputs. In mammalian central nervous system (CNS) neurons, local postsynaptic protein synthesis in dendrites is well established. In contrast, until recently, most evidence for local presynaptic protein synthesis in axons and nerve terminals came from invertebrates and the mammalian peripheral nervous system (Alvarez, et al., 2000). Evidence for presynaptic protein synthesis in the mammalian brain has been difficult to demonstrate largely due to the difficulties of accessing and imaging CNS presynaptic terminals (Akins, et al., 2009). Despite these issues, presynaptic ribosomes have recently been shown to be present in GABA-ergic interneurons in mature mouse hippocampal neurons, where presynaptic protein synthesis is necessary to induce a long-term depression of synaptic responses (Younts, et al., 2016). Local protein synthesis has also been shown to occur in the axons of developing mammalian brain neurons, and plays a role in establishing nerve terminals (Batista, et al., 2017) and affecting release at recently formed nerve terminals (Taylor, et al., 2013). Although it is still highly debated, recent work provides good evidence that local protein synthesis can occur in nerve terminals in mammalian brains and it can affect presynaptic activity.

To better understand the role of presynaptic protein synthesis in the brain, we have used the calyx of Held synapse, located in the medial nucleus of the trapezoid body (MNTB) in the auditory brainstem (von Gersdorff and Borst, 2002). This synapse is involved in sound localization, and can maintain prolonged synaptic transmission at frequencies of 100 to 200 Hz. The calyx of Held is a large, glutamatergic nerve terminal that forms a monosynaptic, axosomatic connection onto MNTB principle cells. This large presynaptic terminal contains hundreds of individual release sites, and the size of the terminal facilitates imaging (Rodriguez-Contreras, et al., 2008; Korber, et al., 2015). In addition, the basic mechanisms of pre- and postsynaptic responses have been extensively characterized at this synapse (Neher, 2017). This synapse also undergoes significant developmental changes in its morphology and physiological characteristics that involves changes in presynaptic protein content, occurring around the onset of hearing in mice, at approximately postnatal day 10 (Borst and Soria van Hoeve, 2012). Finally, in a mouse brain, the calyx of Held nerve terminal is ~3 mm away from the cell body. This distance could enhance the need for local translation at the nerve terminal. These characteristics make this synapse an excellent model for studying the effects of local presynaptic protein synthesis on synaptic transmission.

Our data show that presynaptic ribosomes are present, functional, and active under non-stimulated conditions. In addition, we show that within ~1 hour of inhibiting protein synthesis, there is an increase in the frequency of spontaneous neurotransmitter release, an increase in the paired pulse ratio, and an increased amount of release throughout 100 Hz and 200 Hz stimulus trains. These findings demonstrate that local presynaptic protein synthesis occurs at the calyx of Held nerve terminal, and it affects basal levels of spontaneous neurotransmitter release as well as release during prolonged levels of evoked activity. This represents a previously unknown role for ongoing local translation in adjusting spontaneous and evoked vesicle release.

## Results

### Evidence for presynaptic ribosomes at the calyx of Held nerve terminal

The calyx of Held is a large glutamatergic, monosynaptic nerve terminal located in the medial nucleus of the trapezoid body (MNTB) in the mammalian auditory brainstem (Figure 1A). Cell bodies in the anterior ventral cochlear nucleus project axons a significant distance to the MNTB, which is ~3 mm in a mouse brain (Figure 1A). Up to approximately postnatal day (P) 12, the calyx primarily has a spherical or spoon shaped morphology (Figure 1B, left panels). This large spherical morphology provides a well-defined image of the presynaptic compartment that allows the ability to clearly distinguish fluorescent signals in the presynaptic terminal from fluorescence in the postsynaptic soma. By P12, the calyx terminal begins a change to a fenestrated morphology that is prevalent by P16, when is considered to be mature in morphology and function (Grande and Wang, 2011) (Figure 1B, right panels). The morphological changes are accompanied by changes in protein expression that allow faster action potential kinetics (Yang and Wang, 2006) and synaptic release properties (Borst and Soria van Hoeve, 2012) that begin at ~P10, and allow this synapse to function at the high frequency and fidelity (Taschenberger and von Gersdorff, 2000) that is required for sound localization (Oertel, 1999; Carr, et al., 2001). Accordingly, there is a high level of protein turnover that occurs slightly before and throughout this period.

**Figure 1.**
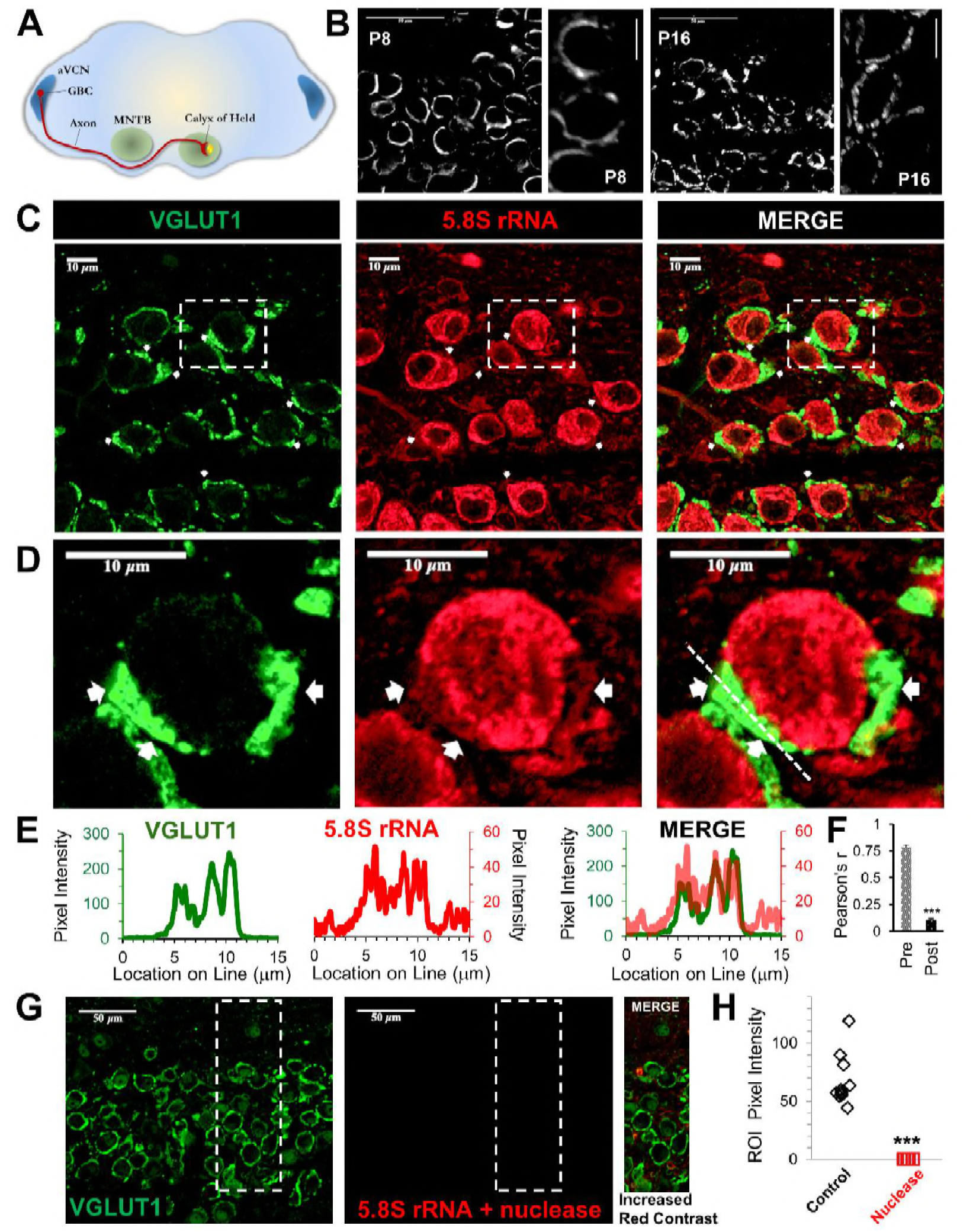
Presence of presynaptic ribosomes in brain slices. **A**. Globular bushy cells (GBCs) in anteroventral cochlear nucleus (aVCN) project long axons (~3mm) forming monosynaptic calyx synapses onto principal cells in contralateral medial nucleus of trapezoid body (MNTB). **B**. VGLUT1 antibody labels presynaptic calyx terminals. At postnatal day 8 (P8) the calyx largely surrounds the postsynaptic neuron. By P16, a fenestrated morphology with swellings is seen (scale: 50 μm or 10 μm). **C**. VGLUT1 antibody labels presynaptic terminals. Immunolabeling of 5.8S rRNA identifies ribosomes. White arrows in each panel depict some examples of clear presynaptic 5.8S rRNA signal. **D**. High optical zoom of dotted box in panel C. White arrows mark presynaptic 5.8S rRNA. **E**. Line scan analysis depicts pixel intensity of VGLUT1 and 5.8S rRNA along a 15 μm line shown in Figure 1D. Merged line scans show excellent overlap in relative signal intensity of VGLUT1 and 5.8S rRNA. **F**. Pearson’s correlation coefficients (r) quantifies colocalization between the VGLUT1 and 5.8S rRNA signals, (n = 10 presynaptic terminals). The r-value of 0.78 ± 0.02, demonstrates high colocalization. The r-values for the matching postsynaptic cell body regions (n = 10) have an r-value of 0.11 ± 0.10 (p ≤ 0.001). **G**. Nuclease treatment prior to 5.8S rRNA antibody binding eliminates the ribosomal signal. Enhanced contrast further shows the lack of ribosomal RNA after nuclease treatment. **H**. Nuclease treatment effectively eliminates 5.8S rRNA signal compared to control conditions (p ≤ 0.001).

We hypothesized that local protein synthesis could occur at this nerve terminal, particularly due to the long axon length, its size (Borst, et al., 1995) and high frequency firing (Wu and Kelly, 1993; Borst, et al., 1995; Taschenberger and von Gersdorff, 2000) that requires high levels of protein to maintain activity at the >600 release sites in a calyx nerve terminal (Satzler, et al., 2002; Taschenberger, et al., 2002; Wimmer, et al., 2006; Dondzillo, et al., 2010). To investigate presynaptic protein synthesis, we first determined whether 5.8S ribosomal RNA (rRNA), a major ribosomal component which is required to execute ribosomal translocation (Lerner, et al., 1981; Abou Elela and Nazar, 1997; Koenig, et al., 2000), is present in the nerve terminal. This component has been shown to be present in dendritic compartments, axons (Koenig, et al., 2000; Zheng, et al., 2001; Spillane, et al., 2013; Taylor, et al., 2013) and neurites, as evidence for the presence of ribosomes (Bolognani, et al., 2004; Kim, et al., 2005; Kim and Kim, 2006; Oyang, et al., 2011). Recent work, using super-resolution microscopy, has shown that 5.8S rRNA is present in nerve terminals of CA1 inhibitory interneuron (Younts, et al., 2016). Given the small size of most presynaptic terminals, standard imaging techniques can be difficult. Therefore, the large size of the calyx of Held nerve terminal helps to determine the presence and localization of 5.8S rRNA in the nerve terminal.

To label the large calyx of Held nerve terminal, an antibody against the vesicular glutamate transporter, VGLUT1, was used (Figure 1C, D; left panel, green), which is standard to label this presynaptic terminal (Billups, 2005; Rodriguez-Contreras, et al., 2008; Fioravante, et al., 2011; Chen, et al., 2013; Kempf, et al., 2013). Immunolabelling for 5.8S rRNA in brain slices shows a robust signal (Figure 1C, D; center panel, red), particularly in neuronal somata, consistent with the high levels of protein synthesis that occur in the cell body (Palay and Palade, 1955; Giuditta, et al., 2008). We typically observe several areas that exhibit clear 5.8s rRNA labelling in the presynaptic terminal, as shown in the representative images (Figure 1C, D). The average intensity ratio for the presynaptic terminal to background signal is 3:1, which allowed us to clearly distinguish the presynaptic signal. As expected, the average intensity for the signal in the presynaptic terminal was less than that of the somata, with an average intensity ratio of 4:1 for somata to presynaptic signals. Despite the strong postsynaptic fluorescence, we were able to unambiguously identify numerous areas with clear presynaptic signals (see white arrows in Figure 1C and D). To determine the overlap of the 5.8S signal with the VGLUT1 signal, we performed line scan analysis (Figure 1E) to compare the positional overlap of VGLUT1 and 5.8S rRNA intensities at higher magnification (Figure 1D; line shown in right panel). We find a high correlation between the relative intensities of the 5.8S rRNA and VGLUT1 signals (Figure 1E, merge), with similar peaks and troughs in their intensities. The troughs in signal intensity could be due to the presence of organelles and other presynaptic components that reduce both signals. Finally, we performed correlation-coefficient analysis to quantify the overlap between VGLUT1 and 5.8S rRNA. We calculated a Pearson’s correlation-coefficient (r) of 0.78 ± 0.04 SEM for VGLUT1 and 5.8S rRNA signal in the presynaptic terminal. For comparison, and to control for the possibility of background fluorescence, we calculated a Pearson’s r-value of 0.11 ± 0.01 SEM in the region surrounded by the VGLUT1 signal, corresponding to the somatic compartment of the postsynaptic neuron (Figure 1F, n = 13 neuronal pairs, p<0.001). This provides very strong evidence that ribosomes are present in the calyx of Held presynaptic nerve terminal.

To verify the specificity of the 5.8S rRNA signal, we treated the brain slices with nucleases to degrade the 5.8S rRNA prior to antibody labelling and found this eliminated the presynaptic and postsynaptic 5.8S signal (Figure 1G, center). To better visualize any residual 5.8S signal remaining after nuclease treatment, we maximized the signal contrast (Figure 1G, right panel), but still found a complete lack of 5.8S signal in nerve terminals and somata. The average pixel intensity for the combined pre- and postsynaptic compartments in untreated slices (68.30 ± 7.33 SEM, n = 10) was far greater than the low remaining signal in nuclease treated slices (0.12 ± 0.01 SEM, n = 10) from the same brain (Fig 1H). These data further demonstrate the presence of ribosomes in the presynaptic terminal, suggesting the ability for local protein synthesis at this nerve terminal.

### Functional presynaptic ribosomes

Our data show that a major ribosomal component is present in the presynaptic terminal. To determine if these ribosomes are fully assembled and functional, we used the <u>SU</u>rface <u>SE</u>nsing of <u>T</u>ranslation (SUnSET) technique. This technique allows us to directly visualize locations of protein synthesis using a fluorescent signal that is proportional to the amount of translation. Briefly, this method uses puromycin, which mimics tRNA and becomes incorporated into nascent polypeptide chains. Specific antibody labeling is then used to detect the amount and location of puromycylation events (Schmidt, et al., 2009; Goodman, et al., 2012; Goodman and Hornberger, 2013). As described below, our results using this technique confirm that functional ribosomes are present in the presynaptic terminal.

A 10 minute application of puromycin allowed us to detect ribosome activity in brain slices. We found fluorescent signal in calyx of Held nerve terminals and in principle cell somata, with relatively low background activity from other nearby cells (Figure 2A,D,E). To quantify the presence of active ribosomes in the presynaptic compartment, we calculated the Pearson’s correlation coefficient (r) for the puromycin and VGLUT1 signals. We find r-values of (0.74 ± 0.05 SEM) for the presynaptic terminal compared to (0.14 ± 0.03 SEM) for the somatic region surrounded by the calyx nerve terminal (Figure 2B, n = 13 neurons, p<0.001). We also find a puromycin signal at some presynaptic terminals where the signal from the postsynaptic neuron is low or absent, likely due to damage or disintegration of the postsynaptic neuron during slice preparation (Figure 2C). Localizing puromycin fluorescence to the presynaptic terminal demonstrates the presence of active presynaptic ribosomes. As expected, the fluorescence intensity is higher in somata than in the terminal. At a higher magnification, puromycin labelling is clearly visible in the presynaptic terminal demonstrating the presence of functional presynaptic ribosomes (Figure 2D,E). To better visualize the relationship between puromycin labelling and the presynaptic marker VGLUT1, we used line scans to assess the degree of colocalization (Figure 2D,E, graphs). We find excellent agreement in the location and relative intensity of the two signals (Figure 2D,E, graphs showing merged line scan overlay), thus demonstrating that translationally competent ribosomes are found in the calyx of Held presynaptic nerve terminal, and they are active under resting conditions. We note that longer puromycin application periods were tested, and this produced a strong pre- and postsynaptic signal, but longer application times also sharply increased the background signal, presumably due to activity from other neurons and glia in the slice (data not shown). We conclude that all of the necessary components (mRNA, tRNA, rRNA, ribosomal proteins and ribosomal binding proteins), which are required to execute translation, must also be located in the nerve terminal.

**Figure 2.**
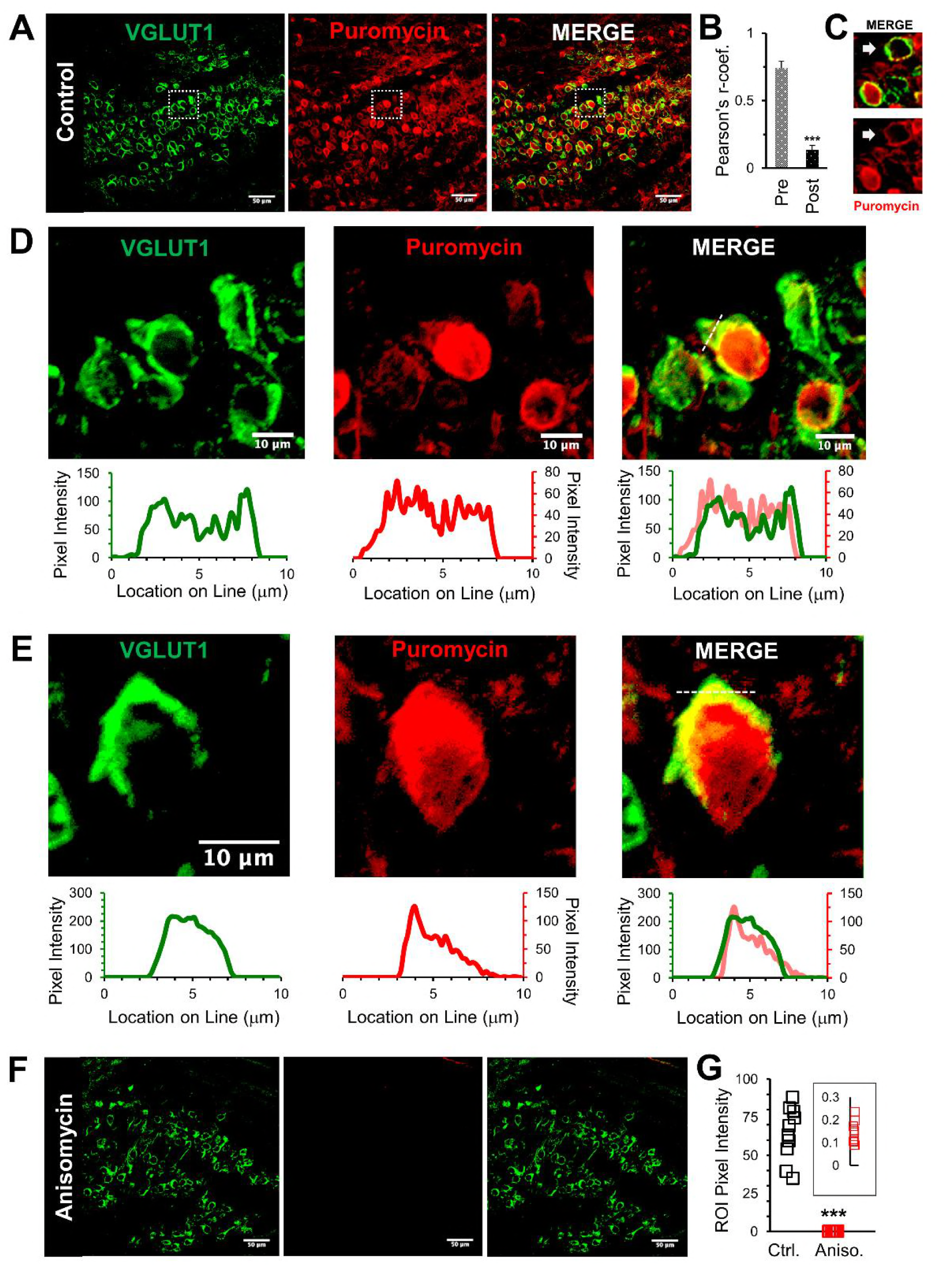
Active ribosomes in the presynaptic terminal in brain slices. **A**. VGLUT1 antibody labels presynaptic terminals. Puromycin, tRNA analog, labelling shows active translation sites (SUnSET). Merge VGLUT1 + Puromycin shows active presynaptic ribosomes. **B**. Pearson’s correlation coefficient (r) quantifies colocalization between VGLUT1 and puromycin signals for presynaptic terminals (n = 10). Average r =0.74 ± 0.05, demonstrates high colocalization. A comparison, average r in corresponding postsynaptic cell bodies =0.14 ± 0.03 (p ≤ 0.001; n = 10). **C**. Example of presynaptic puromycin signal (arrow) in the absence of a postsynaptic signal. Likely due to damage and deterioration of postsynaptic neuron with intact terminal. **D**. Top Panels: higher optical zoom (63x) of dotted box in panel 2A. Bottom Panels: Line scan analysis shows pixel intensity of VGLUT1 and puromycin along the 10 mm line shown in panel 2E. Line scan overlay shows a high overlap in relative signal intensity of VGLUT1 and puromycin along scan line. **E**. Additional example from a different brain slice (PN8) shows active ribosomes in both the pre- and postsynaptic compartments, demonstrated in line scans below each image. Merged overlay shows an excellent match in relative intensities of VGLUT1 and puromycin. **F**. Application of translational inhibitor (anisomycin, 40 μM) for 1 hour prior to puromycin treatment eliminates puromycin labelling (center panel and right panel), showing specificity of puromycin binding to active ribosomes. **G**. Anisomycin treatment eliminates the puromycin signal. Inset shows higher detail of the residual ROI pixel intensity following anisomycin treatment (p ≤ 0.001).

To validate that the SUnSET assay detects active translation, we treated brain slices with the translational inhibitor anisomycin for 1 hour prior to the addition of puromycin (Figure 2F). Ribosomes must be active for puromycin to be incorporated into a polypeptide chain to be detected by the SUnSET assay. Consistent with this, we find that anisomycin treatment effectively eliminates the puromycin signal (Figure 2F, center), giving a >100 fold reduction in the fluorescence intensity following anisomycin treatment (Figure 2G). These results demonstrate that the SUnSET assay provides an efficient and specific measurement of the presence and general location of active ribosomes. This verifies that the necessary components for protein synthesis are present in the presynaptic terminal and are capable of forming translationally competent ribosomes that are active.

### Spontaneous synaptic events indicate presynaptic effects of inhibiting translation

To determine if ongoing protein synthesis affects synaptic transmission, we first looked at miniature excitatory postsynaptic currents (mEPSCs) to determine if spontaneous release events are affected after protein synthesis is inhibited. The frequency of spontaneous events is due to presynaptic release properties (Kavalali, 2015) while the amplitude and shape of the response are largely attributed to postsynaptic changes in ionotropic receptor responses. However, presynaptic properties such as the level of neurotransmitter filling in vesicles can also affect the amplitude of the postsynaptic current (Goh, et al., 2011). Since the need for protein synthesis could be affected by prior activity, it was important to measure spontaneous activity at several times during our recordings to determine if inhibiting translation affects the initial mEPSCs, and the mEPSCs that occur after evoked activity. As noted in the methods section, application of the protein synthesis inhibitor in the physiology experiments, and the subsequent data analysis, were performed blinded.

We find that the initial mEPSC frequency, measured shortly after onset of whole-cell recording, is higher in cells treated with the protein synthesis inhibitor anisomycin (2.4 ± 0.7 Hz, n = 11 recordings) compared to untreated neurons (1.4 ± 0.3 Hz, n = 10 recordings) from the same animals (Figure 3A). The initial cumulative probability histogram of the time between mEPSC events clearly shows that inhibiting protein synthesis decreases the time between mEPSC events compared to control recordings of neurons in untreated slices from the same animals (Figure 3B, p < 0.001, Kolmogorov-Smirnov test). For example, in neurons treated with anisomycin, 70 percent of the events occur with an interval less than ~350 msec (Figure 3B, red dotted line), compared to ~700 msec for control recordings (Figure 3B, black dotted line). To further examine this, we fit exponential curves to the cumulative probability data to quantify the time course of spontaneous release. We find that both the control and the protein synthesis inhibited cumulative probability distributions are well fit by double exponential curves, indicating a fast and slow component of spontaneous release (Figure 3B, grey lines through data points). Both the control and anisomycin treated neurons have a fast component with a time constant of ~170 msec, and a slower component which is 4 to 5 fold slower (Figure 3B). Interestingly, the fast component accounts for the majority of mEPSC events for anisomycin treated neurons (64%), in contrast to control neurons where the fast component only accounts for 37% of the mEPSC event intervals. Thus, the percentage of fast versus slow spontaneous release events were equal but opposite in their distribution, demonstrating a similarity in fast and slow time course but a major difference in the percentage of fast versus slow release events. Therefore, inhibiting protein synthesis has an effect at the presynaptic terminal that causes an increase in the prevalence of fast spontaneous release events.

**Figure 3.**
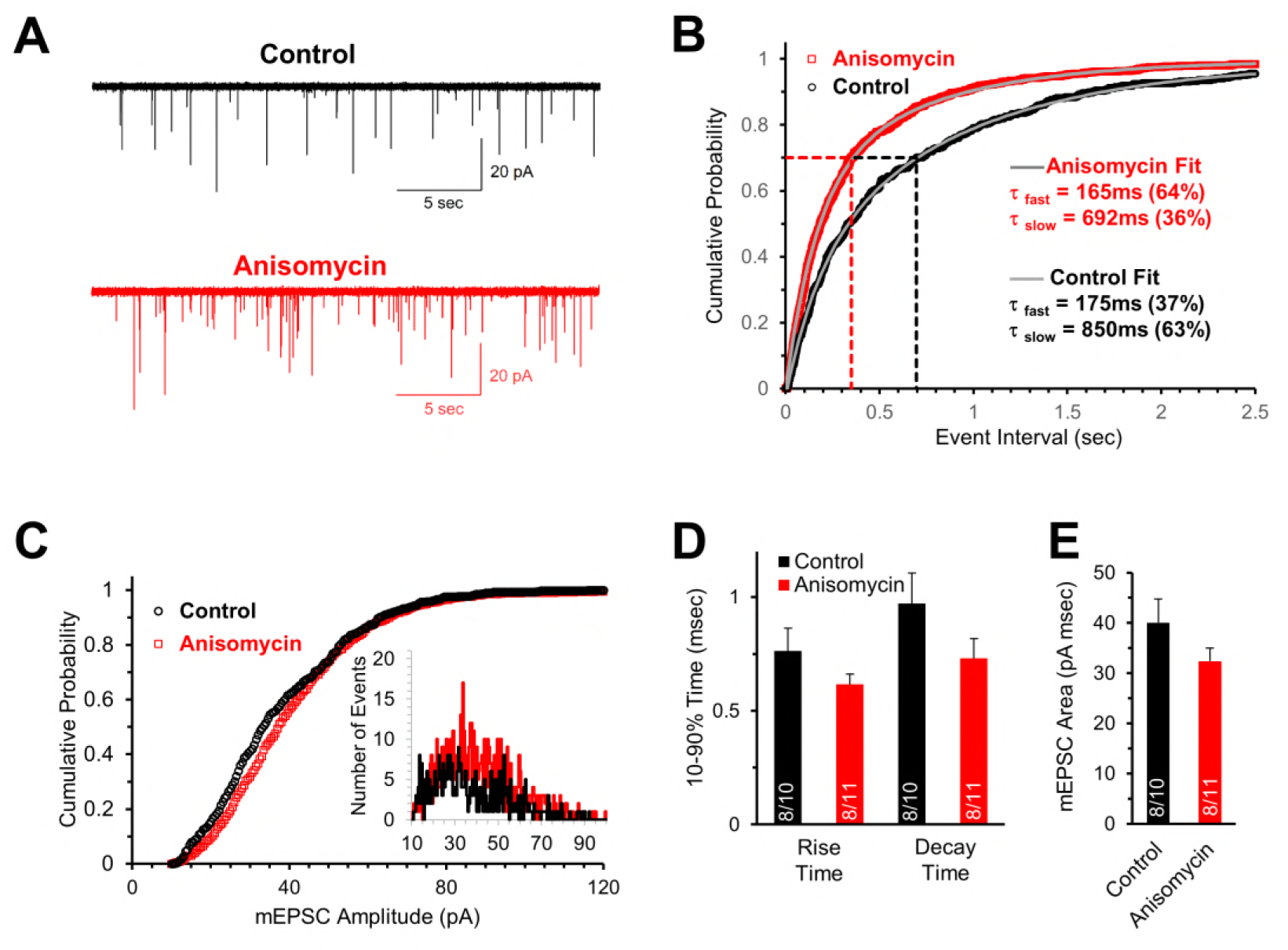
Initial spontaneous release frequency is higher in neurons treated with protein synthesis inhibitor, demonstrating a presynaptic effect. **A**. Representative recordings of the initial spontaneous release in control neurons (black) and neurons treated with the translational inhibitor anisomycin (red). **B**. Cumulative probability of the intervals between spontaneous release events measured by mEPSCs in neurons treated with protein synthesis inhibitor (red) and control neurons (black). The time constants and amplitudes of double exponential fits (gray versus blue lines) show a reversal in the percent contributions of the fast and slow release events in control and translation inhibited neurons. **C**. Cumulative probability of mEPSC amplitudes in protein synthesis inhibited (red) and control (black) neurons. Inset histogram: number of release events at different amplitudes. **D**. Average mEPSC 10-90% rise time (left) and decay time (right) for control (black) and neurons treated with anisomycin (red). (n-values in bar graph: # of animals/ # of neurons) **E**. Average mEPSC area for control (black) and neurons treated with anisomycin (red).

In contrast to the differences seen in the frequency of mEPSCs, the amplitudes of mEPSCs were similar for control (36.8 ± 2.5 pA, n = 10) and protein synthesis inhibited neurons (38.7 ± 2.0 pA, n = 11), as shown in the cumulative probability of the mEPSC amplitudes (Figure 3C) and the amplitude histogram (Figure 3C, inset). For the shape of the mEPSCs, the protein synthesis inhibited neurons have a slightly faster rise time (Figure 3D, p = 0.19), a slightly faster decay time (Figure 3D, p = 0.14), and a slightly smaller average mEPSC area (Figure 3E, p = 0.16) but none of these differences were statistically significant. In summary, the absence of an effect on the mEPSC amplitude, and small non-significant effects on the shape of the mEPSC, demonstrate that the postsynaptic response is not significantly affected (within ~1 hour) by inhibiting protein synthesis. However, the differences in the mEPSC frequency demonstrate that inhibiting protein synthesis has a presynaptic effect on the probability of spontaneous release.

### Enhanced spontaneous release following tetanus eliminates mEPSC frequency differences between control and protein synthesis inhibited neurons

Prolonged stimulation produces a transient elevation in the frequency of spontaneous release events (Habets and Borst, 2006). Given our finding that inhibiting protein synthesis also increases the frequency of spontaneous release events, we determined if the two effects act independently. Accordingly, we measured the frequency of spontaneous release in control and protein synthesis inhibited neurons, before and shortly after a tetanic stimulation at 100 Hz for 4 sec. It is important to note that prior to the pre-tetanus mEPSC recording, the neurons had received several rounds of evoked activity which is discussed in the next section. Although there was a ≥2 minute period without evoked stimulation to allow recovery for the pre-tetanus recording (Fig 4A_1_), there is still a small increase in the mEPSC frequency in both control (2.5 ± 0.45 Hz) and protein synthesis inhibited neurons (3.6 ± 0.89 Hz) compared to the spontaneous frequency measured before any evoked responses were given (Fig 3A). The difference in the timing of mEPSC events in control and protein synthesis inhibited neurons prior to tetanic stimulation is still clearly visible in the cumulative probability histogram of mEPSC event intervals (Figure 4B1; p<0.001, Kolmogorov-Smirnov test). Furthermore, both the protein synthesis inhibited and control neurons continue to show a fast and slow process for spontaneous release, with the fast component accounting for the majority of release event intervals in protein synthesis inhibited neurons (τ_fast_ = 143 msec, 70%; τ_slow_ = 512 msec) and the minority of release event intervals in control neurons (τ_fast_ = 143 msec, 27%; τ_slow_ = 512 msec). Next, we delivered a tetanic stimulation (100 Hz, 4 sec) and measured the mEPSC frequency starting <5 sec after tetanic stimulation. Interestingly, following tetanic stimulation, the frequency of spontaneous release is nearly identical for both control (6.1 ± 0.86 Hz) and protein synthesis inhibited (6.9 ± 1.2 Hz) neurons (Figure 4A2). This is best shown in the cumulative probability histogram of the mEPSC intervals, where the control and protein synthesis inhibited mEPSC curves partially overlap, and no longer have a statistically significant difference (Figure 4B_2_, p = 0.35, Kolmogorov-Smirnov test). Finally, although the fast and slow components of spontaneous release are still present following tetanic stimulation (Figure 4B2), the fast component of spontaneous release sped up (τ ≅ 75-100 msec) and the slow component also became faster (τ ≅ 250-300 msec) following tetanic stimulation for both control and protein synthesis inhibited neurons. Furthermore, following tetanic stimulation, the fast component accounted for approximately 70% of the frequency of spontaneous release events for both control and protein synthesis inhibited neurons. The finding that the cumulative probability of the release intervals overlap following tetanic stimulation demonstrates that the effects of tetanic stimulation are greater for the control conditions. The smaller relative effect of tetanic stimulation after anisomycin treatment could indicate that protein synthesis inhibition and tetanic stimulation have similar presynaptic mechanisms that act to increase the percentage of fast spontaneous release events.

**Figure 4.**
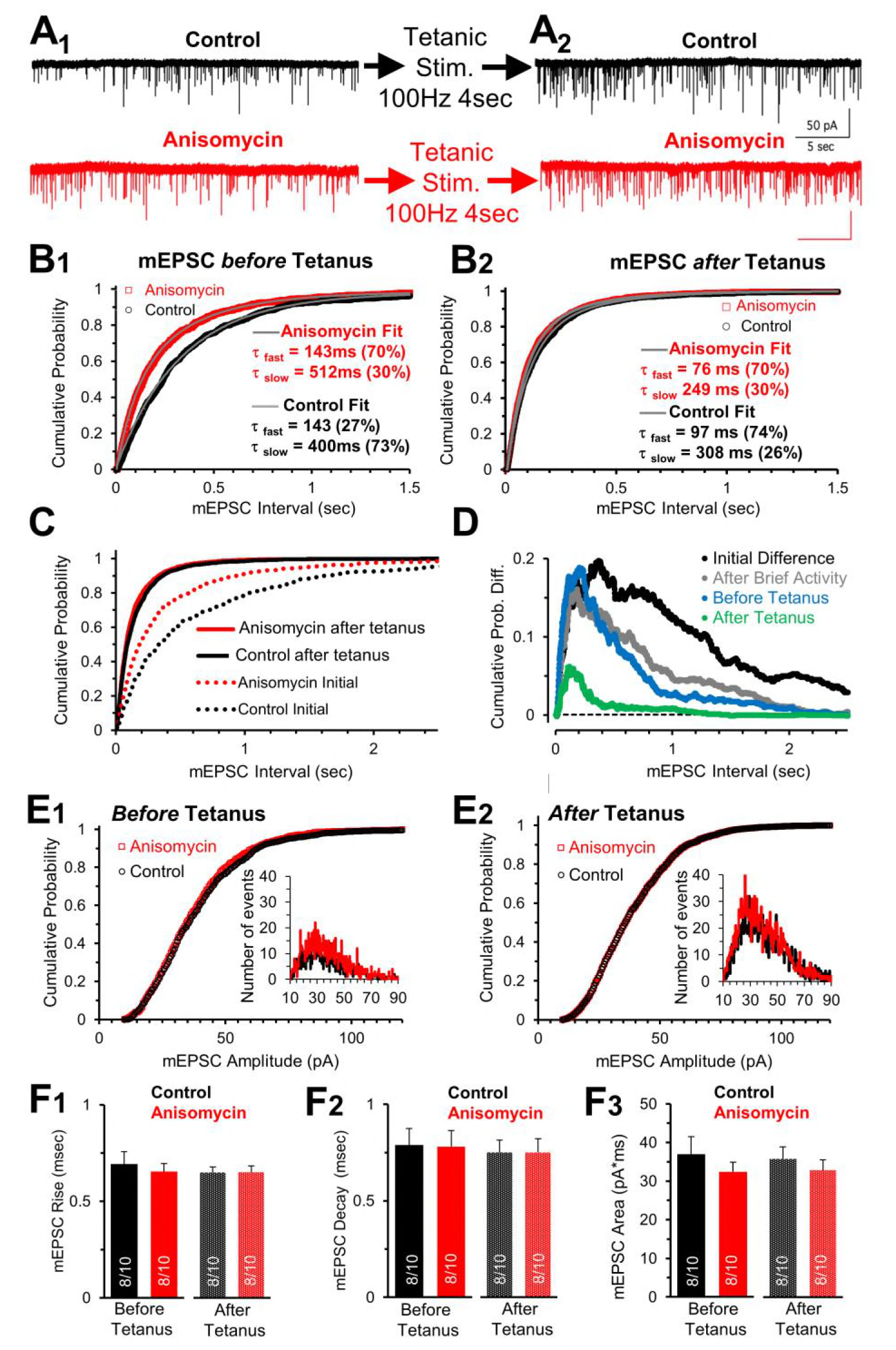
Spontaneous release following tetanus eliminates differences in mEPSC frequency between control and protein synthesis inhibited neurons. **^**A**_1_.^** Spontaneous release in control (black) and slices treated with the translational inhibitor anisomycin (red). Earlier activity (0.4 sec, 100 and 200 Hz, and 4 sec 100 Hz) increased frequency in both groups. ^**A_2_.**^ Spontaneous release in control (black traces) and neurons treated with the protein synthesis inhibitor anisomycin (red) following tetanic stimuli. **^B_1_.^** Cumulative probability of interval times between spontaneous release events in neurons treated with anisomycin (red) and control neurons (black) before tetanic stimulation. Double exponential fits (gray) show differences in the percent contributions of the fast and slow components. ^**B_2_.**^ Cumulative probability of intervals between mEPSC events after a tetanic stimulation. Double exponential fits (gray) show percent differences in the fast and slow components. **C**. Initial cumulative probability of release (dotted lines) compared to release following tetanic stimulation in control (black) and neurons treated with anisomycin. **D**. Differences in cumulative probability of release in anisomycin treated minus control neurons: initial difference (black); after brief activity (100 and 200 Hz for 400 msec each, gray); before tetanic stim. (blue); following tetanic stimulation (green). **E_1_.** Cumulative probability of mEPSC amplitudes in protein synthesis inhibited (red) and control (black) neurons prior to evoked stimulation. Inset: depicts histogram of the mEPSC amplitudes. **E_2_.** Cumulative probability of mEPSC amplitudes in protein synthesis inhibited neurons (red) and control (black) following a tetanic stimulation. Inset: mEPSC amplitude histogram. **F_1_.** mEPSC 10-90% rise time for control (black) and anisomycin (red) before (left) and following tetanic stimulation (right). (n values in bar graph apply to all panels in the figure: # of animals / # of neurons) **F_2_.** Average mEPSC 10-90% decay time for control (black) and neurons treated with anisomycin (red) before (left) and following tetanic stimulation (right). **F_3_.** Average mEPSC area for control (black) and neurons treated with anisomycin (red) before (left) and following tetanic stimulation (right).

Comparing the initial mEPSC intervals present at the onset of each recording, before evoked stimulation (Figure 3B), with the intervals following tetanic stimulation (Figure 4B_2_) shows that tetanic stimulation increases the fast and slow rates of release, and the percentage of rapid spontaneous release events (Figure 4C). To further show how spontaneous release differs between protein synthesis inhibited neurons and control neurons, we took the initial cumulative probability distributions (Figure 3B) and subtracted the average values in control neurons from the average values in the anisomycin treated neurons (Figure 4D, black trace). We also show how activity affects this difference. Previous activity reduces the difference in the distribution of the release events between protein synthesis inhibited and control neurons (Figure 4D, black versus grey lines), and a tetanic stimulation nearly eliminates these differences (Figure 4D, blue versus green lines). Therefore, the presynaptic effects of evoked activity act to speed up the rate of spontaneous release in control and protein synthesis inhibited neurons. In addition, a tetanic stimulation further increases the rate of release for control and inhibited neurons, and temporarily alters the control neurons so that the majority of spontaneous events occur by the fast component of release.

In contrast to the changes in spontaneous release, the amplitudes of the mEPSCs were unaffected by anisomycin treatment, tetanic stimulation or both combined (Figure 4E_1_ and E_2_). Furthermore, the mEPSC rise times (Figure 4F_1_), decay times (Figure 4F_2_), and area (Figure 4F_3_) were unchanged by tetanic stimulation, inhibiting protein synthesis, or by both combined. Collectively, these results show that inhibiting protein synthesis and presynaptic tetanic stimulation both have little to no effect on the amplitude and shape of the postsynaptic currents generated by spontaneous release. Therefore the effects of inhibiting protein synthesis, and the effects of tetanic stimulation on mEPSC properties are specific to presynaptic effects on the rate of spontaneous release.

### Amplitude and shape of initial evoked responses are relatively unaffected after inhibiting protein synthesis

The finding that spontaneous release of synaptic vesicles is affected by inhibiting protein synthesis suggests that evoked responses could also be affected. In our initial tests, we stimulated at a low frequency (0.1 Hz) to determine if the peak amplitude, shape, and latency of evoked excitatory postsynaptic currents (EPSCs) are affected when protein synthesis is inhibited. Consistent with the results from the mEPSC measurements, the shape of the EPSCs appear to be unaffected by inhibiting protein synthesis over the time course of 45 to 120 minutes of inhibition (Figure 5A). The 10-90% rise and decay times for control and protein synthesis inhibited neurons were similar, with small differences that were not significant (Figure 5A–C). In addition, the average peak amplitudes of the initial responses were also similar for control (5.7 ± 0.51 nA) and protein synthesis inhibited neurons (5.37 ± 0.73 nA), with no statistically significant difference (Figure 5D, p = 0.6). Consistent with this, the average area of the initial EPSCs were also relatively unaffected between treated and control neurons (Figure 5E, p = 0.23). Finally, we also measured the latency of the EPSC responses and found similar values for control and treated neurons (Figure 5F, p = 0.27). Taken together, these data indicate that the kinetics and peak amplitude of the initial evoked postsynaptic responses are relatively unaffected by inhibiting translation over a 1 to 2 hour time course, which is similar to our finding that mEPSC amplitude and shape are not affected by inhibiting translation. However, given the change in spontaneous release events, we anticipated a change in evoked release events which was not apparent when we measured individual EPSCs at the onset of our recordings. To further pursue this, we next tested if inhibiting protein synthesis affects high frequency EPSC responses.

**Figure 5.**
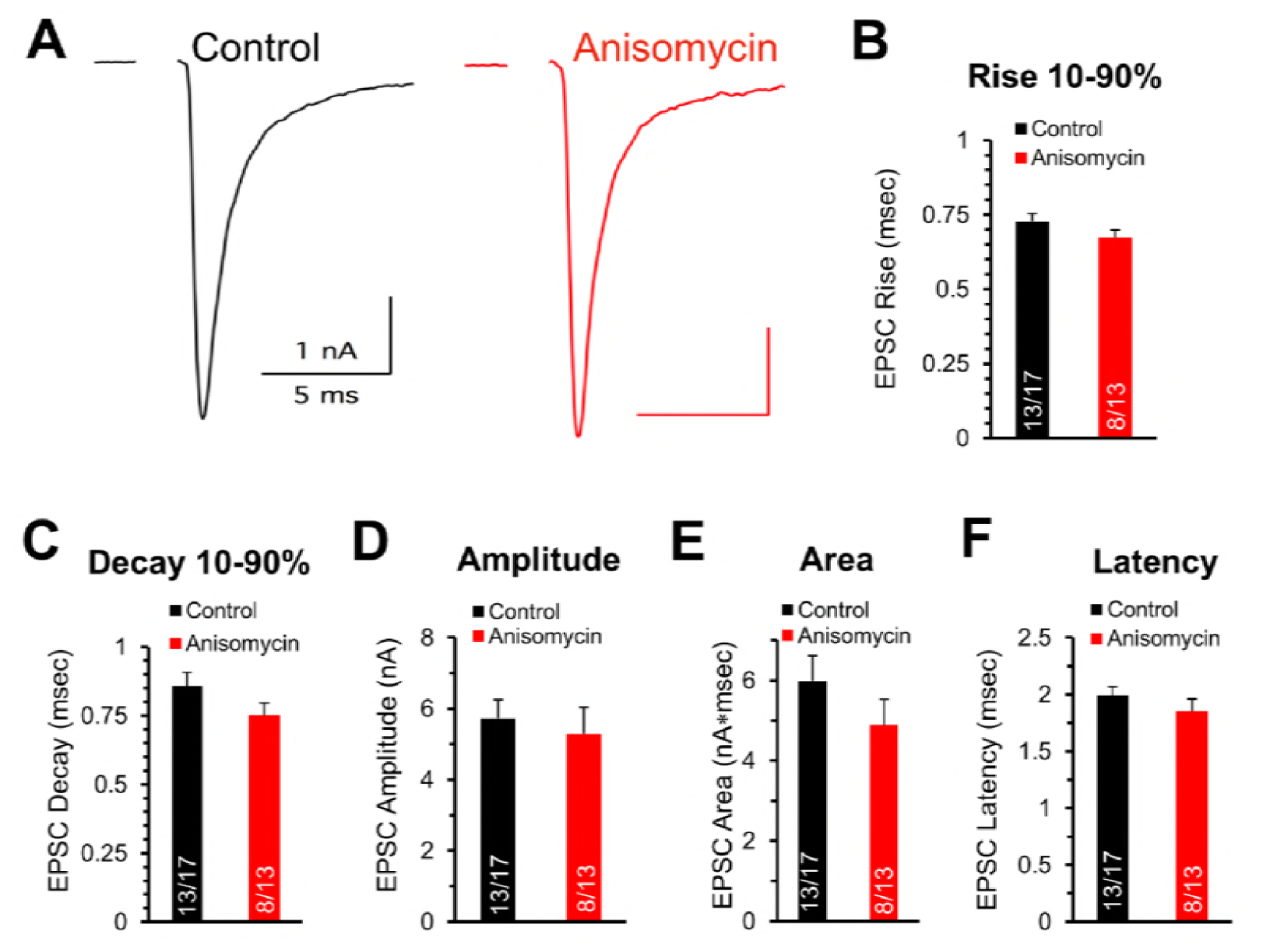
Initial evoked response amplitude and shape are not affected by inhibiting protein synthesis. **A**. Representative traces of evoked excitatory postsynaptic currents (EPSCs) for control (black) and neurons treated with anisomycin (red) from PN10 mice. **B**. Average 10-90% evoked EPSC rise time for control (black) and neurons treated with anisomycin (red). The n-values in bar graphs presented as # of animals/ # of neurons. **C**. Average 10-90% evoked EPSC decay time for control (black) and neurons treated with anisomycin (red). **D**. Average evoked EPSC amplitude, control (black) and anisomycin treated neurons (red). **E**. Average evoked EPSC area, control (black) and neurons treated with anisomycin (red). **F**. Average evoked EPSC latency for control (black) and anisomycin treated (red).

### Paired pulse measurements indicate a presynaptic effect of translational inhibition

The calyx of Held can fire at high frequencies with a high level of precision. We hypothesized that presynaptic protein synthesis may play a role in presynaptic mechanisms of synaptic transmission. Although the amplitude and shape of the initial evoked responses do not appear to be significantly affected by inhibiting protein synthesis (Figure 5), high frequency stimulation recruits additional presynaptic components that could be affected by inhibiting protein synthesis. Therefore, we tested short stimulus trains at 200 Hz for 400 msec which were followed two or more minutes later by a stimulus at 100 Hz. At both frequencies, we observed a tendency for lower levels of depression in the responses from translation inhibited neurons. To quantify this, we first measured the effects of high frequency stimulation on the paired pulse ratio at 200 Hz, measured as the second EPSC (P2) response divided by first EPSC (P1) response (Figure 6A_1_ and A_2_). We find paired pulse depression at an interpulse pulse interval (IPI) of 5 msec in control cells (0.72 ± 0.07 SEM, n = 9 cells from 7 animals) but a slight facilitation in translation inhibited cells at the same interval (1.09 ± 0.09 SEM, p = 0.004, n = 8 cells from 6 animals). To determine if this ratio was affected following prolonged activity, we also measured the 5 msec IPI paired pulse ratio > 2 minutes after a tetanic stimulation (100 Hz for 4 sec). The paired pulse ratio was slightly decreased following tetanic stimulation, which eliminated the small facilitation seen in the protein synthesis inhibited cells (Figure 6A_2_). Next, we compared EPSCs in control and protein synthesis inhibited conditions throughout a 400 msec train at 200 Hz stimulation (Figure 6B_1_). Consistent with the paired pulse results, we observe a reduction in EPSC depression throughout the train in the protein synthesis inhibited responses compared to controls. Accordingly, we directly compared the average responses for the entire train and found that the reduced depression seen after inhibiting protein synthesis occurred throughout the 400 msec train at 200 Hz (Figure 6B_2_). In contrast, for the blinded experiments in which DMSO was applied, the treated and untreated responses were very similar (Supplementary Fig. 1A). Therefore, inhibiting protein synthesis results in higher levels of neurotransmitter release that can be sustained (80 EPSCs) even at a brief 5 msec IPI.

**Figure 6.**
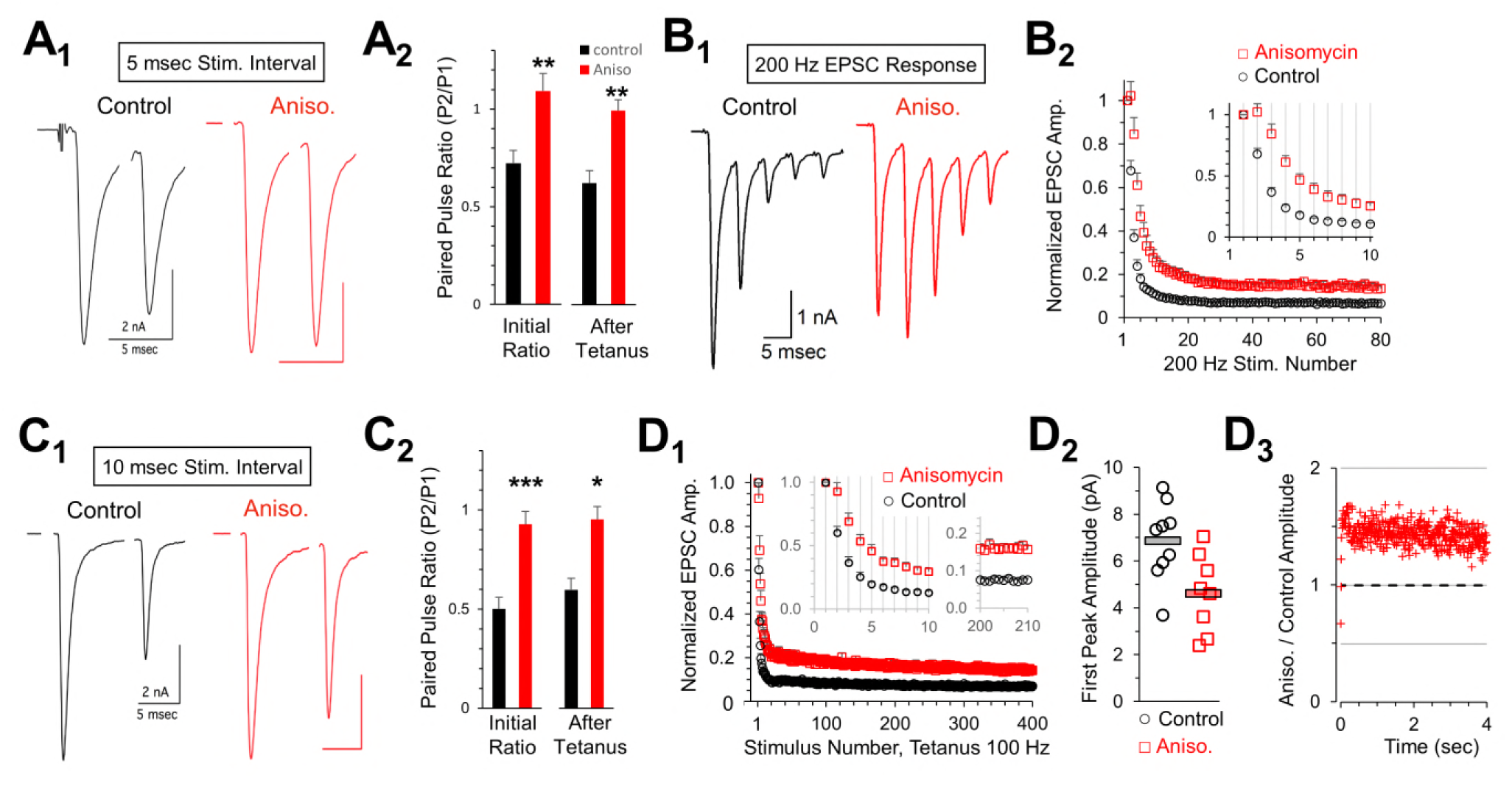
Reduced depression at 100 or 200 Hz after protein synthesis inhibition. **A_1_.** Representative traces of initial paired pulse responses, 5 msec interpulse interval (IPI). **A_2_.** Initial paired pulse ratio (5 msec IPI) for control (black) and anisomycin treatment (red), compared to ratio after tetanic stimulation (100Hz, 4 sec), control (black), anisomycin (red). **B_1_**. Representative EPSC responses at 200 Hz stimulation for control and anisomycin. **B_2_.** Average normalized EPSCs at 200 Hz stimulation for control and anisomycin treatment. **C_1_.** Representative initial paired pulse responses at a 10 msec interpulse interval (IPI). **C_2_.** Initial paired pulse ratio (10 msec IPI) for control (black) and anisomycin treatment (red), compared to ratio (10 msec IPI) after tetanic stimulation of 100Hz for 4 sec. **D_1_.** Average normalized EPSCs from tetanic stimulation in control and anisomycin treated neurons. **D_2_.** Amplitudes of first EPSC in tetanic stimulation response for each control neuron (black circles) and anisomycin treated neuron (red squares). Horizontal bars corresponds to the average responses. **D_3_.** Ratio of 100 Hz EPSC responses from anisomycin and control neurons shows anisomycin treated neurons maintain a higher EPSC responses during sustained stimulation for 4 sec

Facilitation and depression are affected by the interval between pulses. Accordingly, we also measured the paired pulse ratio at 10 msec IPI (Figure 6C_1_ and C_2_). At this interval, additional depression occurred in control recordings (0.50 ± 0.06 SEM, n = 8 cells from 7 animals) but protein synthesis inhibited responses exhibited only weak depression (0.93 ± 0.06 SEM, n = 8 cells from 7 animals; p<0.001). We also tested how prior exposure to prolonged stimulation (4 sec at 100 Hz) affects the paired pulse ratio and found a slight increase in control neurons (0.60 ± 0.07 SEM, n = 8 cells), but no change in the protein synthesis inhibited responses (0.95 ± 0.11 SEM, n = 8 cells). This indicates that the lack of paired pulse depression in protein synthesis inhibited synaptic responses is stable. In summary, the differences we observe in the paired pulse depression at 5 and 10 msec IPI are consistent with differences in presynaptic mechanisms involving vesicle release (von Gersdorff and Borst, 2002; Fioravante and Regehr, 2011), indicating that inhibiting translation has an effect on vesicle release that reduces paired pulse depression.

To determine how the slight paired pulse depression at 10 msec IPI affects responses throughout prolonged trains, we measured all EPSPs in response to a 4 sec tetanic stimulation at 10 msec IPI (100 Hz). Similar to the results at 5 msec IPI (200 Hz), we find inhibiting protein synthesis reduced depression throughout the entire duration of the 4 sec stimulation at 100 Hz compared to control responses (Figure 6D_1_). This effect was not seen when DMSO was applied during the blinded experiments (Supplementary Fig. 1B). Due to variability in the amplitudes of EPSCs in different recordings, it is necessary to normalize the EPSC responses to the peak amplitude of the first response. In addition, facilitation or reduced paired pulse depression often occurs when the initial probability of release is reduced (Fioravante and Regehr, 2011), although other mechanisms may exist (Neher, 2017). To determine if normalization affected our results, we compared the average P1 amplitude for each cell tested, and found a slight reduction in the average P1 amplitude in protein synthesis inhibited responses (4.61 ± 0.59 SEM) compared to control (6.85 ± 0.59 SEM; p = 0.015; Figure 6D_2_). Compared to the amplitudes at the onset of the recordings (Figure 5D) this represents a small increase in control and decrease in protein synthesis inhibited responses. This difference suggested that normalizing the data could affect the results. Therefore, we directly compared the ratio of the non-normalized peak responses in protein synthesis inhibited conditions divided by control conditions (Figure 6D_3_). Aside from the first three responses, the peak amplitudes in the protein synthesis inhibited responses was ~1.4 fold higher than the peak amplitudes of control responses. This demonstrates that translation inhibited cells maintain a higher amount of release, due to lower depression, compared to responses from control cells even during prolonged stimulation for 4 sec (Figure 6E) indicating that this resistance to depression is robust. While it is completely possible, that the effects of inhibiting protein synthesis could change with repeated activity occurring over several hours, or be accompanied by other changes in synaptic responses, we are unable to reliably measure this given the need to maintain long recordings with consistent control responses over longer time periods. Therefore the enhanced synaptic response may be an initial consequence of inhibiting protein synthesis, and different or additional effects are likely to occur over hours or days.

### Inhibiting protein synthesis affects vesicle release and recycling

The presynaptic effects on paired pulse ratios and increased response levels that occur throughout trains of prolonged stimulation suggest that the readily release pool (RRP), vesicle release, and vesicle recycling could be affected by inhibiting protein synthesis. To measure this, we graphed the cumulative EPSC response during a 100 Hz stimulation train (Figure 7A_1_). A best fit line through the EPSC responses to stimuli 15 to 30 (Figure 7A_1_) provides the slope of the beginning of the steady state response, and the y-intercept of this line provides a measurement of the readily releasable pool (Schneggenburger, et al., 1999; Neher, 2015). Using this method, the average size of the RRP (Figure 7A_2_) was the same for control neurons (16.3 ±1.6 nA, n = 9 cells from 7 animals) and protein synthesis inhibited neurons (anisomycin, 15.97 ± 1.9 nA, n = 8 from 7 animals; p = 0.89). This indicates that the initial capacity for vesicular release of neurotransmitter is not changed by inhibiting protein synthesis. Next, the initial probability of release was measured by dividing the peak amplitude the first EPSC response by the corresponding RRP measured for that train (Schneggenburger, et al., 1999). Here, we find the initial release probability (Pr) for control neurons (0.43 ± 0.03) is higher than the value in anisomycin treated neurons (0.29 ± 0.02; p = 0.003), therefore the differences we see in the initial peak response in protein synthesis inhibited neurons is explained by a reduction in the initial probability of release (Figure 7A_3_). Lastly, the slope of the steady state of the cumulative response (Figure 7A_1_) provides a measurement of the rate of vesicle recycling. We find that the vesicle recycling rate (Figure 7A_4_) increases when protein synthesis is inhibited (96.9 ± 1.6 pA/msec), compared to the rate in control neurons (62.2 ± 3.5 pA/msec; p = 0.037). This indicates that an increased rate of vesicle replenishment is responsible for the increased responses following anisomycin treatment (Figure 6B_2_, D_1_, and D_3_).

**Figure 7.**
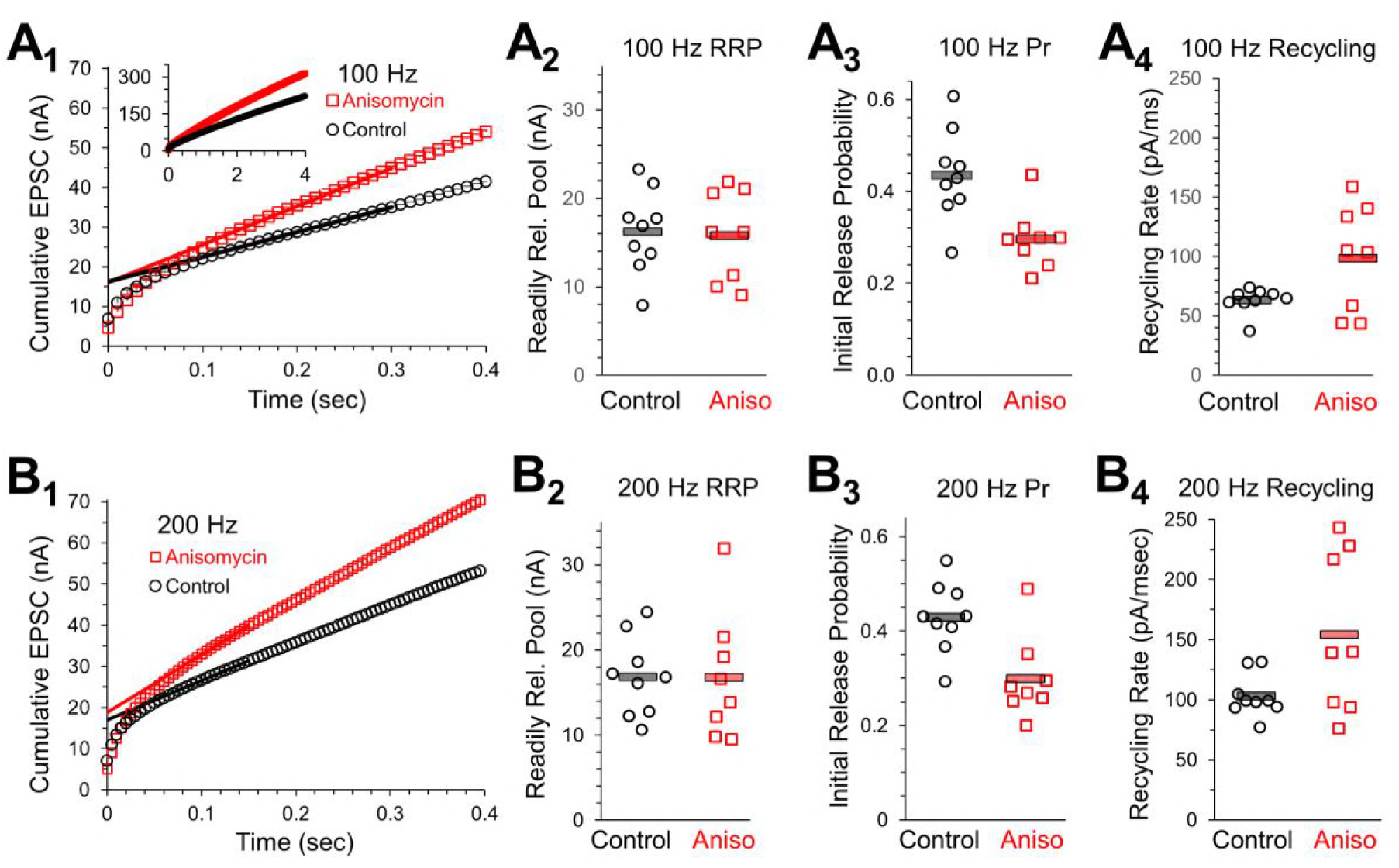
Initial release probability and vesicle recycling are affected by inhibiting protein synthesis. **A_1_.** 100 Hz Cumulative EPSC plot of averaged peak response in control (black) and protein synthesis inhibited (anisomycin, red) neurons. The y-intercept of a line fit to responses 20 to 30 measures the readily releasable pool (RRP). **A_2_.** The initial probability of release (Pr_1_) measured during 100 Hz trains, measured as the ratio of the amplitude of the first EPSC to the RRP. Averages shown by horizontal bars. **A_3_.** RRP measurement for each cell, during 100 Hz stim, measured as the y-intercept of a best fit line through responses 20 to 30, for the beginning of the steady state response. **A_4_.** Recycling rate for steady state response during 100 Hz stimulation, measured as the slope of a best fit line through responses 20 to 30, separately measured for each cell. **B_1_.** 200 Hz Cumulative EPSC plot of averaged peak response in control and protein synthesis inhibited (Anisomycin) neurons. The y-intercept of a line fit to responses 20 to 30 measures the readily releasable pool (RRP). **B_2_.** The initial probability of release (Pr_1_) measured during 200 Hz trains, measured as the ratio of the amplitude of the first EPSC to the RRP. Averages shown by horizontal bars. **B_3_.** RRP measurement for each cell, during 200 Hz stim, measured as the y-intercept of a best fit line through responses 20 to 30, for the beginning of the steady state response. **B_4_.** Recycling rate for steady state response during 200 Hz stimulation, measured as the slope of a best fit line through responses 20 to 30, separately measured for each cell.

To provide additional estimates of vesicle properties, we also measured responses from stimulation at 200 Hz (Figure 7B_1_). As anticipated, we found that the readily releasable pool values for control and protein synthesis inhibited neurons are the same at 100 Hz and 200 Hz (Figure 7A_2_ and 7B_2_). At 200 Hz, the RRP value for control neurons is 16.8 ± 1.5 nA compared to 16.8 ± 2.6 nA for anisomycin treated neurons (p > 0.9). As expected, the initial probability of release at 200 Hz is nearly identical to the values measured at 100 Hz stimulation (Figure 7A_3_), with a Pr of 0.43 ± 0.02 for control and 0.3 ± 0.03 for anisomycin treated neurons (p = 0.004) estimated from the EPSC responses generated by 200 Hz stimulation (Figure 7 B_3_). However, the vesicle recycling rate, as measured by the slope of the steady state response in the cumulative EPSC graph, is higher at 200 Hz compared to 100 Hz (Figure 7A_4_ and 7B_4_), as expected given the need to maintain release levels. At 200 Hz, the vesicle recycling rate in control neurons (102.8 ± 5.9 pA/msec) was similar to the vesicle recycling rate of anisomycin treated neurons at 100 Hz (96.9 ± 1.6 pA/msec; p = 0.72). Furthermore, at 200 Hz, the anisomycin treated neurons maintained a faster recycling rate (154.1 ± 23.4 pA/msec) compared to control neurons (p = 0.04). Interestingly, the magnitude of the increase in the vesicle recycling rate from 100 to 200 Hz stimulation is nearly identical for control (1.65) and anisomycin treated neurons (1.59). The finding that the vesicle recycling rate increase that occurs from 100 to 200 Hz is nearly the same in control and anisomycin treated neurons demonstrates that the factors that scale up the recycling rate are not affected by inhibiting protein synthesis. Instead, it appears that the normal steady state levels of release and recycling are increased after protein synthesis is inhibited.

## Discussion

We hypothesized that local translation occurs in the presynaptic compartment and that it is necessary to maintain normal levels of neurotransmitter release. In support of this, we have shown that 5.8s rRNA, a major component of ribosomes, is present in the presynaptic terminal at the calyx of Held synapse. This provides further evidence that presynaptic ribosomal components are present in established mammalian CNS nerve terminals (Younts, et al., 2016) in addition to developing neurites and axons (Taylor, et al., 2013; Batista, et al., 2017). We verified that presynaptic ribosomes are functional, using the SUnSET technique which has previously been used to demonstrate local translation in dendritic, axonal and neuritic compartments (Batista, et al., 2017) in cell cultures. Due to the large size of the calyx of Held we were able to employ this technique in mammalian brain slice to show that presynaptic ribosomes are present and functional, producing a presynaptic signal within tens of minutes. Notably, this signal showing active translation at functional ribosomes is completely blocked when brain slices are pretreated with a translational inhibitor. These data provide convincing evidence that local protein synthesis occurs at the calyx of Held nerve terminal.

In addition to demonstrating active translation in presynaptic ribosomes in the Calyx of Held, we find that ongoing protein synthesis affects neurotransmitter release. Starting with spontaneous activity, we found that inhibiting translation causes an increase in the frequency of spontaneous release. Specifically, the spontaneous release events show a fast and slow component in the intervals between events, and inhibiting translation increases the percentage of fast release events to ~65% of the spontaneous release intervals. In control neurons we found the opposite relationship, where the slow release events account for ~65% of the event intervals. Therefore, ongoing protein synthesis limits spontaneous release by favoring slow release events. The finding that ongoing protein synthesis acts to lower the spontaneous release frequency is consistent with the idea that the frequency of spontaneous release is highly controlled, given that spontaneous events function as important signals in synaptic development and homeostasis (Kavalali, 2015). For example, spontaneous release of glutamate can suppress local protein synthesis in dendrites (Sutton, et al., 2006). Furthermore, NMDA receptors that are specifically activated by spontaneous release appear to be responsible for the rapid antidepressant effect produced by ketamine exposure (Autry, et al., 2011). Therefore, over the last decade, accumulating evidence demonstrates that spontaneous release can have significant effects on synaptic function. In addition, vesicles that undergo spontaneous release may preferentially come from a population separate from the vesicles that respond to evoked release (Sara, et al., 2005), and this appears to involve association with specific proteins (Hua, et al., 2011). Finally, vesicles involved in spontaneous release can have different sensitivity to intracellular calcium than vesicles involved in evoked release, and spontaneous release may be able to occur independent of intracellular calcium (Schneggenburger and Rosenmund, 2015). Therefore vesicles that undergo spontaneous release appear to be controlled separately from vesicles that fuse in response to action potentials. Spontaneous release is a highly regulated process that is important in maintaining synaptic function, and we show that ongoing protein synthesis plays a role in limiting the frequency of spontaneous release events.

Tetanic stimulation has previously been shown to transiently increase the frequency of spontaneous release, due to residual calcium (Korogod, et al., 2005) and may also involve PKC activation (Korogod, et al., 2007; Fioravante, et al., 2014). In our work, we show that increased frequency of spontaneous release following tetanic stimulation is due to an increase in the time course and percentage of the fast spontaneous release events in control neurons. Despite having initially opposite levels of fast and slow spontaneous release, control and protein synthesis inhibited responses have matching percentages of fast and slow release events following a tetanic stimulation. Accordingly, the increased spontaneous release that occurs after inhibiting protein synthesis may in part involve similar mechanisms that increase spontaneous release after tetanic stimulation.

In addition to the effects on spontaneous release, we also observe facilitation or reduced depression in paired pulse ratios when protein synthesis is inhibited. Interestingly, although we found that the readily releasable pools are identical in protein synthesis inhibited and control conditions, the initial probability of release during a train of stimulation is lower when protein synthesis is inhibited. However, during prolonged stimulation, a reduced amount of depression is maintained throughout the stimulation. Furthermore, we also find an increase in the vesicle recycling rate when protein synthesis is inhibited, indicating that the higher levels of release require a faster recycling rate to maintain the elevated amount of release (Sara, et al., 2002; Qiu, et al., 2015). Therefore, despite the reduced probability of release for the first response in the stimulation trains, the overall levels of release and subsequent vesicle recycling are elevated after inhibiting protein synthesis.

Importantly, the shape and amplitude of spontaneous and evoked responses are unaffected during the 1-2 hours that protein synthesis is inhibited in our experiments, indicating that the postsynaptic receptor responses are not affected. The lack of change in the postsynaptic membrane currents also argues against a possible retrograde signal that is affected by inhibiting protein synthesis. In support of this, in experiments where the postsynaptic neuron was lysed and removed, presynaptic release properties were not changed (He, et al., 2006). Therefore, the differences in synaptic responses under control and protein synthesis inhibited conditions are due to presynaptic effects on neurotransmitter release. The effect on spontaneous and evoked neurotransmitter release properties, combined with the presence of active presynaptic ribosomes, indicate that presynaptic proteins which act to limit synaptic transmission can be synthesized locally in the presynaptic terminal.

The ability to synthesize some proteins locally makes particular sense in cells that have long processes, such as axons and dendrites. The transient rate of axoplasmic transport has been reported to be ~0.1 to 1 μm/sec, and sustained rates are slower (Maday, et al., 2014). In addition, translation at the cell body requires retrograde axonal transport of a signal from the nerve terminal to the cell body, followed by subsequent production of protein and the anterograde axonal transport of protein for delivery to the nerve terminal. Therefore, local synthesis of at least some regulatory proteins saves significant time, allowing a local neuronal region to rapidly upregulate some essential proteins in response to changes in neuronal activity. While significant work has been done over the last two decades to demonstrate the presence and properties of local protein synthesis in dendritic compartments of CNS neurons (Rodriguez, et al., 2008; Rangaraju, et al., 2017), evidence demonstrating the presence and requirements for local presynaptic protein synthesis in intact CNS mammalian neurons is very recent (Younts, et al., 2016). However, work using mammalian synaptosomes has produced evidence of mRNA transcripts and the ability to generate newly synthesized proteins (Alvarez, et al., 2000), although potential contamination from postsynaptic neurons or glia has been a major concern. Additional evidence comes from mRNA found in axons (Alvarez, et al., 2000) and recently formed nerve terminals (Batista, et al., 2017). These presynaptic transcripts code for a variety of proteins including some that can affect vesicle recycling and fusion such as β-catenin, β-tubulin, and β-actin. In addition, transcripts for nuclear encoded mitochondrial proteins have been found in axons. It is important to note that in our experiments, the calyx nerve terminal is no longer connected to the neuronal cell body because the axons are severed during brain slicing. Given the lack of connection between the cell body and nerve terminal, newly synthesized proteins in the calyx nerve terminal cannot come from the cell body that gives rise to the axon that forms the nerve terminal. Based on our imaging data, we conclude that ongoing protein synthesis is occurring in the presynaptic terminal, although some amount could also occur in the adjacent section of the axon.

The work shown here is the first demonstration that local presynaptic translation occurs in established nerve terminals in situ in mammalian brain slices. Our finding that ongoing protein synthesis helps to limit vesicle release has interesting implications for how nerve terminals maintain and modulate their presynaptic release properties. Limiting evoked synaptic responses effectively increases presynaptic efficiency by allowing the nerve terminal to conserve some of the substantial energy involved in vesicular release, retrieval, refilling, and recycling (Rangaraju, et al., 2014Rangaraju, et al., 2014; Shulman, et al., 2015; Sobieski, et al., 2017). Limiting synaptic release should also help the nerve terminal to maintain sufficient responses for a longer time. This indicates an important function for newly synthesized proteins over a time course that doesn’t allow transport from the cell body to the presynaptic terminal. The further study of the presynaptic processes that are affected by inhibiting translation will help us to better understand how spontaneous and evoked release are controlled, and the role that local protein synthesis plays in maintaining and modulating synaptic responses.

## Acknowledgements

We thank Drs. Mark Plummer, Bonnie Firestein, Ulrich Hengst, Zhiping Pang, Daphné Robinson, and Lu-Yang Wang for discussions, critique and suggestions on this work. We also thank Drs. Kelvin Kwan, Noriko Goldsmith, and Wise Young (W. M. Keck Center for Collaborative Neuroscience, Rutgers), Dr. 0Nanci Kane (Waksman Institute, Rutgers), Daniel Martin (Biomedical Engineering, Rutgers), Dr. Lorin Milescu and Dr. Mirela Milescu (University of Missouri) for advice, assistance, and use of imaging systems.

## Ethics

For Animal experimentation, all procedures were carefully performed and were approved by the Rutgers University Institutional Animal Care and Use Committee (IACUC; protocol 10-062).

## Author Contributions

MS designed and performed experiments, performed data analysis, interpreted results and drafted the manuscript. RK performed data analysis, interpreted results, assisted in editing the manuscript. MB performed data analysis, interpreted results and contributed calyx diagram. KP conceived the project, designed and performed experiments, performed data analysis, interpreted results and drafted then edited the manuscript.

## Methods

### Slice Preparation and Electrophysiology

#### Brain slices

C57BL6 mice (Charles River Laboratories) from postnatal day 8 to 12, of either sex were used for all experiments described. The mice were housed in a facility approved by the Association for Assessment and Accreditation of Laboratory Animal Care International, and protocols used for handling and care were reviewed by the Rutgers University Animal Care and Facilities Committee. Animals were decapitated without prior anesthesia, in accordance with NIH guidelines. Transverse brainstem slice thickness varied from 100 μm (immunohistochemistry and imaging) to 180 μm (electrophysiology) and were generated using a Leica VT1200 vibratome. Throughout the process of dissection and slicing, the brain was maintained in a low-calcium artificial CSF (aCSF) solution at 1-2°C containing the following (in mM): 125 NaCl, 25 NaHCO_3_, 2.5 KCl, 1.25 NaH_2_PO_4_, 25 glucose, 0.8 ascorbic acid, 3 myo-inositol, 2 Na-pyruvate, 3MgCl_2_, and 0.1 CaCl_2_, pH 7.4, when oxygenated with carbogen gas (95% oxygen, 5% carbon dioxide). Once produced, slices were transferred to a holding chamber maintained at ~35°C for 30-40 min in normal calcium aCSF solution with the same composition listed above except for 1mM MgCl_2_ and 2mM CaCl_2_. This same solution was also used as the standard recording solution for electrophysiology experiments (see below). All experiments were performed at room temperature (22-25°C) for up to ~4-5h after the recovery period.

#### Electrophysiology

Patch-clamp recordings were conducted using an EPC10 USB double patch-clamp amplifier with PatchMaster software (HEKA; Harvard Bioscience). A transverse slice orientation was used in all postsynaptic voltage-clamp recordings in order to maintain the integrity of the calyceal axons for fiber stimulation. Calyx synapses in the medial nucleus of the trapezoid body (MNTB) were afferently stimulated (A-M Systems Isolated Pulse Stimulator Model 2100) using a bipolar fiber stimulator (lab design) placed at the midline of the slice. The MNTB field was scanned with an extracellular pipette to locate neurons that respond to midline fiber stimulation. For whole-cell recording, patch pipettes were produced from thick-walled borosilicate glass, 2.0 mm outer diameter, 1.16 mm inner diameter (Sutter Instruments). Postsynaptic pipettes (2-3 MΩ) were filled with a solution containing (in mM): 125 Cs-methanesulfonate, 20 CsCl, 20 TEA, 10 HEPES, 5 phosphocreatine (Alpha Aesar), 4 ATP, 0.3 GTP, and 2 QX-314 Cl^−^ (Sigma Aldrich; to block voltage gated Na^+^ channels on the postsynaptic neuron to measure the true EPSC), and was buffered to pH 7.4 using CsOH. Postsynaptic series resistances (R_s_) for voltage clamp recordings were typically 4-10 MΩ. An R_s_ compensation of 75-80% was applied for all recordings such that the adjusted R_s_ was in the range of 2-5 MΩ. Cells for which these criteria could not be applied, or maintained, were excluded from analysis. Recordings were acquired at a sampling frequencies of 20 KHz, and filtered by a 4-pole Bessel filter at 3kHz. Holding potentials were set to −65mV; junction potentials, calculated to be −11 mV, were not corrected.

#### Blinded testing and analysis conditions and criteria for testing anisomycin

For all electrophysiology recordings and data analysis measurements, anisomycin treatment and control conditions were blinded. In addition, the quality of the slices, neurons, and general recording conditions were determined by 1-2 initial recordings in normal aCSF. If the initial recordings had stable responses that lasted the duration of the stimulus protocols, a minimum of 25 minutes, then the recording solution was switched to a blinded cylinder of recording solution for subsequent recordings which were performed after ~ 1 hour of treatment (45 minutes to 120 minutes) in the absence of fiber stimulation. Since spontaneous action potentials are not present in these recording conditions, only spontaneous release activity occurred during the treatment period. On each day, the blinded cylinder would contain either: 40μM anisomycin (Sigma Aldrich) or DMSO alone (vehicle). To test the effect of the translational inhibitor anisomycin (40μM) on synaptic response characteristics, slices were preincubated for ~1hour in the presence of the drug in the absence of afferent fiber stimulation. At all times, aCSF was continuously circulated using a peristaltic pump; total volume of the solution was 30mL. All recordings, control and test conditions, were made in the presence of 25μm bicuculline and 2μm strychnine to block inhibitory responses. Power analysis to determine the appropriate sample size was performed based on means and standard deviation values of preliminary data. Recordings from 5 cells in each condition was estimated to be adequately powered, for α = 0.5, and a 0.8 power of test.

#### Recordings from MNTB neurons and Data Analysis

Miniature excitatory post-synaptic currents (mEPSCs) were recorded during 30 sec continuous recordings at several times during the stimulation protocol. mEPSCs were analyzed by Mini Analysis Software (Synaptosoft). The following mEPSC search parameters were used: gain, 20; blocks, 3,940; threshold, 10 pA; period to search for a local maximum, 20,000 μsec; time before a peak for baseline, 5,000 μsec; period to search a decay time, 5,000; fraction of peak to find a decay time, 0.5; period to average a baseline, 2,000 μsec; area threshold, 10; number of points to average for peak, 3; direction of peak, negative). Analysis was performed using the above settings, and visually checked to ensure accuracy. Evoked response traces were exported to Igor Pro (Wavemetrics, Portland, OR), and measurements were made manually, or using Taro Tools (Igor macro, Taro Ishikawa) with visual inspection and adjustment as necessary for every measured peak amplitude.

Data are presented as mean ± standard error of the mean (SEM). Unless otherwise noted, a Student’s t-test was employed to determine statistical significance. Calculated *p*-values are indicated in relevant figures as follows: *p* ≤ 0.05 is considered significant (*); *p* ≤ 0.01 very significant (**); and *p* ≤ 0.001 highly significant (***). We define biological replicates as each tested cell (number of recordings), and technical replicates as multiple tests on a single cell. In our experiments, a minimum of four recordings, of spontaneous activity, 30 sec each, were made during the recording time. Data were analyzed as initial spontaneous release levels, and spontaneous release following activity as described in the text. Outlier data for spontaneous event recordings resulted in removal of two cells from the data, as determined by Grubb’s test with α = 1%. The two-sample Kolmogorov-Smirnov (KS) test was used to calculate the *p*-value for the cumulative probabilities of the mEPSC event intervals for two different conditions. Briefly, this non-parametric test uses the maximum vertical difference between two cumulative probability graphs and the total number of measurements to determine the statistical significance of the differences between two cumulative probability distributions. Histograms with identical bin-ranges were used to compare the mEPSC intervals for the two different conditions. This calculation was preformed manually, and by the KS function in MATLAB, which gave very similar or identical values.

### Immunohistochemistry and Confocal Microscopy

#### Immunohistochemistry

Either sex of C57BL6 mice, postnatal (PN) day 8 to 12, (n = 18) were decapitated without previous use of anesthesia, and transverse auditory brainstem slices (100-140μm thick) were prepared as described above. Following recovery in normal aCSF, sections were transferred to a 12 well culture plate and washed 2x in phosphate buffered saline (PBS) (in mM: 137 NaCl, 2.7 KCl, 4.3 Na_2_HPO_4_*7H_2_0, and 1.4 KH_2_PO_4_, pH 7.4). Following the washes, the solution was replaced with ice-cold PBS containing 4% (wt/vol) PFA and fixed for 30 min at room temperature, with gentle agitation. After fixation the sections were rinsed 3x with PBS, and incubated in blocking and permeabilization buffer in PBS containing 10% (vol/vol) normal goat serum (MP Biomedicals, LLC), 2% (wt/vol) BSA and 0.25% (vol/vol) Triton X-100 (Alfa Aesar) for 1h30m at room temperature. Slices were again rinsed with PBS 3x, 10 min for each wash. Sections were further blocked in PBS containing 40μg/mL of AffiniPure Fab Fragment Goat Anti-Mouse IgG (H+L) for 1 hour at room temperature. Slices were then washed (3x) and placed in PBS containing the following; 1% (vol/vol) normal goat serum, 1% (vol/vol) BSA, 0.25% (vol/vol) Triton X-100, and mouse monoclonal anti-5.8S rRNA, clone Y10b at 1:500 (Abcam, ab37144) overnight at 4°C. Importantly, to minimize the possibility if non-specific interactions, all double labeling experiments were done sequentially. Following overnight incubation, slices were washed 3x in PBS, and placed in primary antibody solution containing guinea pig polyclonal anti-vesicular glutamate transporter 1 (VGLUT1) at 1:500 (Synaptic Systems) and allowed to incubate overnight at 4°C. Slices were rinsed 3x in PBS, and placed in PBS containing the following; 1% (wt/vol) BSA, 0.05% (vol/vol) Tween-20, and Alexa-594-conjuagted AffiniPure Goat Anti-Mouse IgG (H=L)(1:500) secondary antibody (Jackson, 115-585-003) for 2h at room temperature. Slices were rinsed 3x in PBS and then incubated in the same buffer as above, but with Alexa-488 conjugated AffiniPure Donkey Anti-Guinea Pig IgG (H+L) (1:500) secondary antibody (Jackson, 706-545-148) for 2h at room temperature. Sections were then washed with PBS and mounted to a glass slide, excess PBS was removed, a few drops of Fluoromount™ (Sigma Aldrich) was added, and covered with Gold Seal cover slips #1.5 (Thermo Fisher). Slides were stored at 4°C.

To confirm 5.8S rRNA specificity we pretreated slices with nucleases. Following fixation sections were washed in PBS and incubated in PBS containing 0.25% (vol/vol) Triton –X100 for 45 minutes. Slices were then washed 3x (10m each) in enzyme buffer (50 mM Tris and 5 mM CaCl_2_, pH=8). Next, slices were incubated in enzyme buffer containing 80 μg/mL RNase A (Fermentas, EN0531) and 300 U/mL micrococcal nuclease (New England Biolabs, M0247) for 60m at 37°C.(Note: control experiments were performed by incubating slices in enzyme buffer (with no enzymes) at 37°C) Following incubations, slices were rinsed 3x (10m each) with PBS and blocked for 1h30m in PBS containing 10% (vol/vol) normal goat serum and 2% (wt/vol) BSA, at room temperature. Antibody application and further slice processing is the same as described above.

#### Confocal Microscopy

Confocal image stacks of PFA treated brain slices were acquired using a Leica TCS SP2 laser-scanning microscope (Leica, Heidelberg, Germany). Image acquisition was performed in sequential scanning mode, using a 3 scan average for each image. Images were displayed and analyzed using FIJI. Regions of interest (ROI) were used to calculated image intensity of individual presynaptic terminals, postsynaptic principal cells, or both. Co-localization was performed by selecting neurons from the same-stacked images at random. Analysis was performed on the image where the neuron was widest. To reduce non-specific signal, a background subtraction value of 16 was subtracted from each image. The FIJI plugin JACop120 (just another co-localization plugin) was used for colocalization analysis which generated a graphical output table that contained the Pearson’s correlation coefficient. Line scan analysis to qualitatively assess signal overlap in a given area was performed in FIJI using the line-plot profile feature. In all experiments performed the VGLUT1 signal was always higher in relative intensity than 5.8S rRNA and puromycin.

### SUnSET Labeling of Newly Synthesized Proteins

#### Puromycylation Assay

Transverse brainstem slices were prepared as described above.

To detect newly synthesized proteins, slices were incubated in puromycin (1.8 μM Sigma Aldrich), added to normal aCSF, for 10 minutes following the brain slice recovery period. Control slices were pre-incubated first with 40 μM anisomycin for 60 min, and puromycin was added at the 50 min time point so that anisomycin and puromycin were present for the last ten minutes. Following incubation slices were rinsed 3x with pre-warmed PBS, prior to fixation. The slices were then further processed as described above, post-fixation. To detect puro-polypeptides a mouse monoclonal anti-puromycin antibody, clone 12D10 (EMD Millipore, MABE343) was used at a dilution of 1:250. The secondary antibody used for indirect immunofluorescence was Alexa-594-conjuagted AffiniPure Goat Anti-Mouse IgG (H=L) at a dilution of 1:500. Imaging was carried out as described above.

---

**Figure S1.**
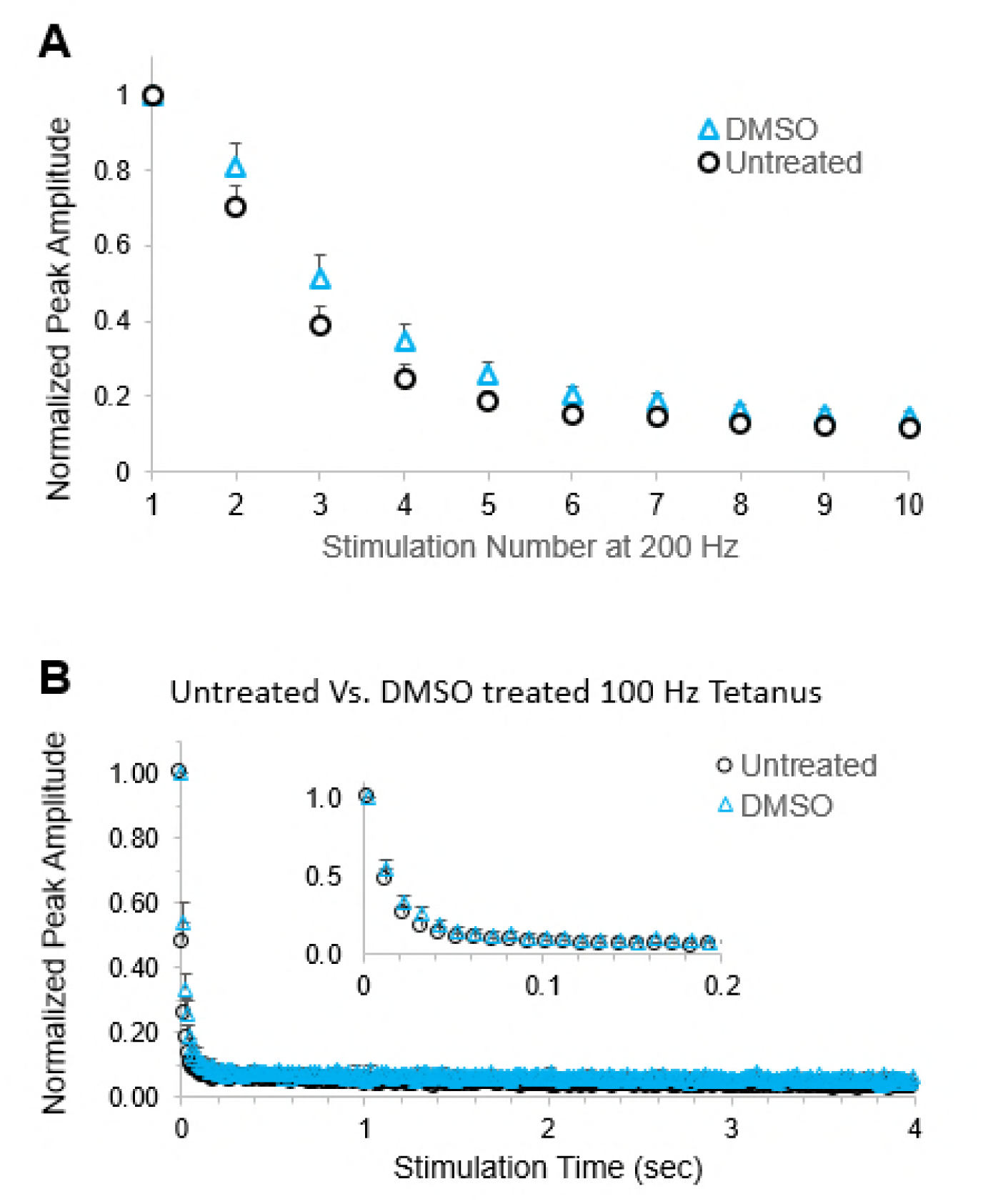
Blinded experiments with DMSO treatment. **A**. Average normalized EPSCs at 200 Hz stimulation for untreated versus DMSO treatment for 1-2 hours (p>0.075). **B**. Average normalized EPSCs at 100 Hz stimulation for untreated versus DMSO treatment for 1-2 hours. No significant difference was found between DMSO and untreated cells (p>0.1).

